# TRPV2 and TRPM2 channels facilitate pulmonary endothelial barrier recovery after ROS-induced permeability

**DOI:** 10.1101/2025.02.13.638083

**Authors:** Lena Schaller, Thomas Gudermann, Alexander Dietrich

## Abstract

Reactive oxygen species (ROS), such as hydrogen peroxide (H_2_O_2_), are known signaling molecules that increase endothelial barrier permeability. In this study, we investigated the roles of two redox-sensitive transient receptor potential (TRP) ion channels, TRPM2 and TRPV2, in H_2_O_2_- induced endothelial barrier dysfunction. Using primary human pulmonary microvascular endothelial cells (HPMEC), we employed impedance-based resistance measurements, Western blot, and immunofluorescence staining to assess the effects of H_2_O_2_ on the endothelial barrier. Exposure to sublytic concentrations of H_2_O_2_ caused an acute loss of endothelial barrier integrity, accompanied by the cleavage of vascular endothelial cadherin (VE-cadherin), which was also apparent after application of the TRPV2 activator cannabidiol. The inhibition of either TRPV2 with tranilast or a disintegrin and metalloprotease domain-containing protein 10 (ADAM10) with GI254023X significantly reduced H_2_O_2_-induced VE-cadherin cleavage, while TRPM2 inhibition by econazole significantly increased H_2_O_2_-driven VE-cadherin cleavage. Although inhibition of either TRP channel did not prevent the initial loss of barrier resistance upon H_2_O_2_ exposure, both were essential for the subsequent recovery of barrier integrity. Time-course immunofluorescence stainings revealed that HPMEC barrier recovery involved a transient localization of N-cadherin proteins at cell-cell junctions, which were replaced by VE-cadherin within 90 minutes. This process of cadherin-switching did not occur upon inhibition of TRPV2 or ADAM10. Our results highlight complementary roles for TRPM2 and TRPV2 as redox sensitive ion channels in the microvascular endothelium and provide insight into the mechanisms underlying pulmonary microvascular endothelial barrier recovery.

**Graphical Abstract: See text for more details:** 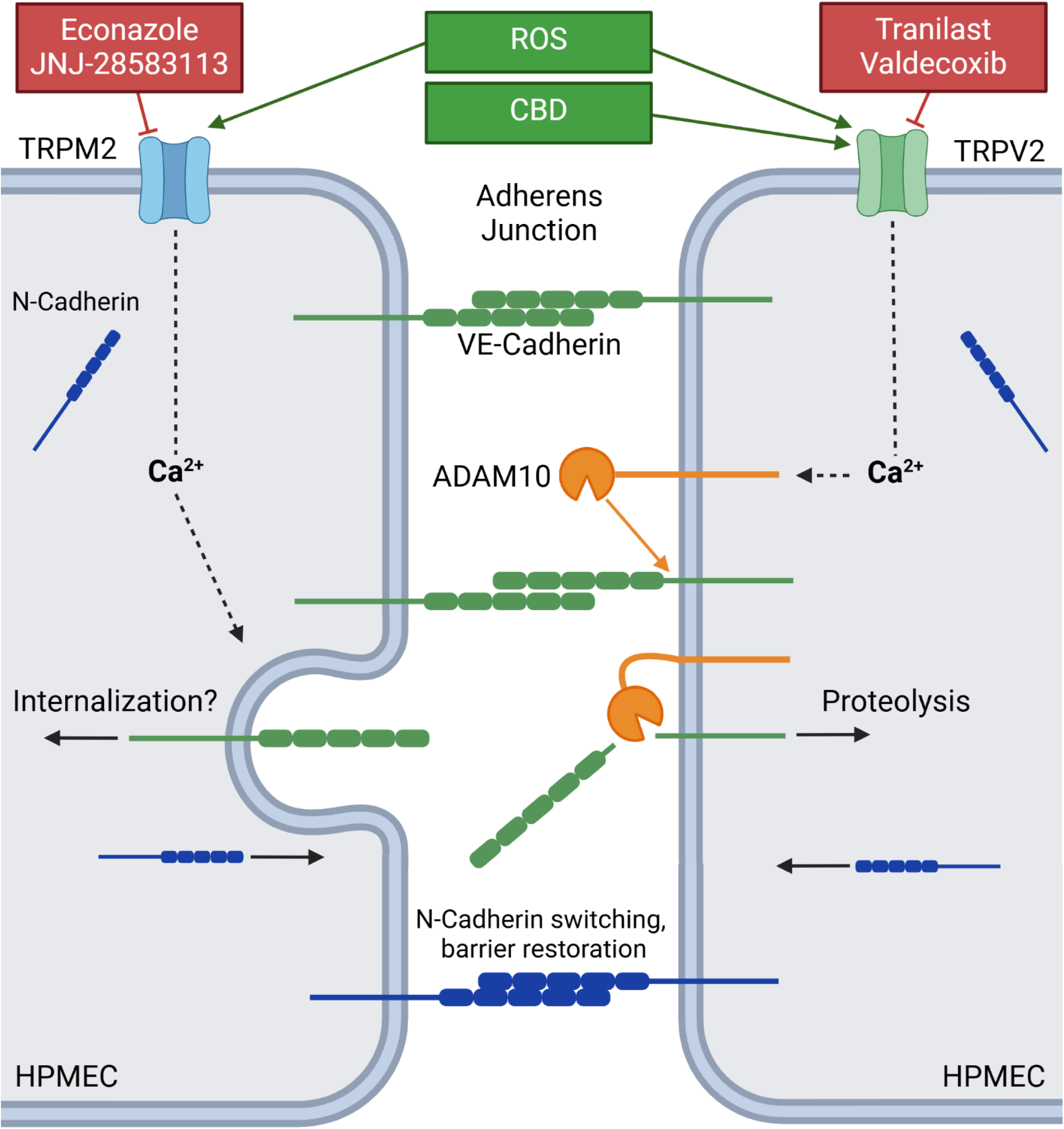

## 1. Introduction

The barrier formed by the pulmonary microvasculature is constitutively restrictive, preventing both pathogen infiltration and edema formation while facilitating the exchange of gases and nutrients between the bloodstream and surrounding tissue [1,2]. While a transient increase in permeability supports biological functions such as wound repair, angiogenesis and immune cell trafficking [3,4], prolonged or extensive permeability can result in pulmonary edema, acute respiratory distress syndrome (ARDS) [5] and atherosclerosis [6].

Reactive oxygen species (ROS), such as hydrogen peroxide (H_2_O_2_), are known effectors of altered endothelial barrier function [7–10]. ROS can arise from exogenous triggers, including infection, ionizing radiation or toxicants, but also occur naturally in the body, such as during mitochondrial respiration [11,12]. The concept of an “oxidative window” describes the optimal range of ROS levels that facilitate cellular processes such as neovascularization, cell proliferation and wound healing [7,8,13]. Deviations from this balance, resulting in oxidative or reductive stress, lead to cellular dysfunction [8].

Adherens junctions (AJs), comprised of Ca^2+^-dependent, homotypic adhesions between the vascular endothelial cadherin (VE-cadherin) proteins of neighboring cells, are essential components of the endothelial barrier [3,14]. While the formation of AJs depends on extracellular Ca^2+^, an increase in intracellular Ca^2+^ can induce endothelial barrier permeability [3,15]. Members of the Transient Receptor Potential (TRP) superfamily form nonselective cation channels that conduct Ca^2+^, and several TRP channels have been implicated in Ca^2+^-induced barrier dysfunction [3]. It has been reported that TRP-induced Ca^2+^ influx could activate a disintegrin and metalloprotease domain-containing protein 10 (ADAM10) [16], a metalloprotease known to cleave VE-cadherin at its extracellular domain [17]. However, a ROS-driven, ADAM10-mediated cleavage of VE-cadherin has yet to be reported in pulmonary microvascular endothelial cells.

TRPM2 is a recognized mediator of ROS-induced Ca^2+^ influx. Expressed in the brain, immune cells, and vasculature, TRPM2 forms a tetrameric, nonselective ion channel that conducts Ca^2+^ and is gated by adenosine diphosphate ribose (ADPR) [18–21], which is generated as a result of ROS-induced DNA damage [18,19]. While TRPM2 is a known modulator of pulmonary endothelial barrier permeability, its knockdown does not completely abolish endothelial Ca^2+^ influx following ROS exposure, suggesting the involvement of additional redox sensitive Ca^2+^ channels [9,22].

The second member of the vanilloid TRP subfamily, TRPV2, is a potential candidate for the undefined source of ROS-induced pulmonary endothelial Ca^2+^ influx. Originally associated with mechanoreception [23], TRPV2 also operates as a redox-sensitive ion channel [24,25], and is highly expressed in the microvascular endothelium in relation to other redox sensitive TRP channels, including TRPM2 and TRPV4 [26]. While TRPV2 has been linked to changes in blood-brain barrier integrity [27], there is no evidence to date linking the channel to altered pulmonary microvascular endothelial barrier function [28].

Here, we applied pharmacological inhibitors to investigate the role of TRPM2 and TRPV2 in H_2_O_2_-induced pulmonary endothelial barrier dysfunction. Neither channel was responsible for the initial loss of barrier resistance, but both channels facilitated the subsequent recovery of barrier integrity. In this model, TRPV2 mediated AJ integrity by inducing ADAM10-driven VE-cadherin cleavage, which was further increased upon TRPM2 inhibition. Endothelial barrier recovery was characterized by the translocation of neural cadherin (N-cadherin) to the plasma membrane upon VE-cadherin removal, suggesting a role for TRP-mediated cadherin switching in the restoration of endothelial barrier function following ROS-induced permeability.

## 2. Methods

### Cells

Primary human pulmonary microvascular endothelial cells (HPMECs) [29] from healthy donors were obtained from Promocell (Heidelberg, Germany, #C-12281) and cultured in endothelial cell growth medium MV (Promocell, #C-22020) at 37 °C and 5 % CO_2_, and were kept until passage 12. Donor information is provided in Supp. Table S1. Relevant ethical statements were provided by Promocell. For experiments involving pharmacological inhibition, HPMECs were pre-incubated for 1 hr in DMEM containing the respective inhibitor(s), which were also present during the subsequent exposure period.

### Quantification of endothelial barrier resistance

HPMECs were seeded onto electrical cell-substrate impedance sensing (ECIS) plates at a density of 8 x 10^4^ cells/well (Applied Biophysics, Troy, NY, USA, 8W10E+), which had been treated with 10 mM of L-Cysteine according to the manufacturer’s recommendation. HPMEC barrier resistance was measured at 4000 Hz using the ECIS ZΦ device (Applied Biophysics), and experiments were conducted once the monolayer resistance had reached a constant state (after ∼48 hr).

## 3. Results

### 3.1. H_2_O_2_ exposure at non-cytolytic concentrations increases HPMEC barrier permeability and triggers ADAM10-dependent VE-cadherin cleavage

Using H_2_O_2_ to mimic ROS production in response to infection, radiation or other toxicants, we monitored changes in barrier resistance of human pulmonary microvascular endothelial cells (HPMEC). While H_2_O_2_ exposure did not exert detectable cytolytic effects after 2 hours (Supp. Fig. S1), changes in HPMEC barrier resistance were observed within 5 minutes of exposure (Fig. 1A). 15 minutes post H_2_O_2_ addition (40 minutes after starting the experiment in Fig 1A), mean HPMEC barrier resistance dropped to 45% ± 11% and 47% ± 14% of the control in cells treated with 75 μM and 300 μM H_2_O_2_, respectively, with recovery noted only in HPMECs treated with 75 μM H_2_O_2_ (Fig. 1B). Additionally, Western blot analysis revealed that H_2_O_2_ exposure caused the formation of a single ∼35 kDa VE-cadherin C-terminal fragment (CTF) (Supp. Fig. S1B, C), the formation of which was prevented by the addition of the specific ADAM10 inhibitor GI254023X [30] (GI254, see Supp. Table S2 for IC_50_ values) (Fig. 1C, D). ADAM10-dependent regulation of VE-cadherin was also observed in immunofluorescence stainings, which showed that ADAM10 inhibition reduced the loss of VE-cadherin signal at cell-cell junctions after H_2_O_2_ exposure (Fig. 1E and F).

**Fig. 1:**
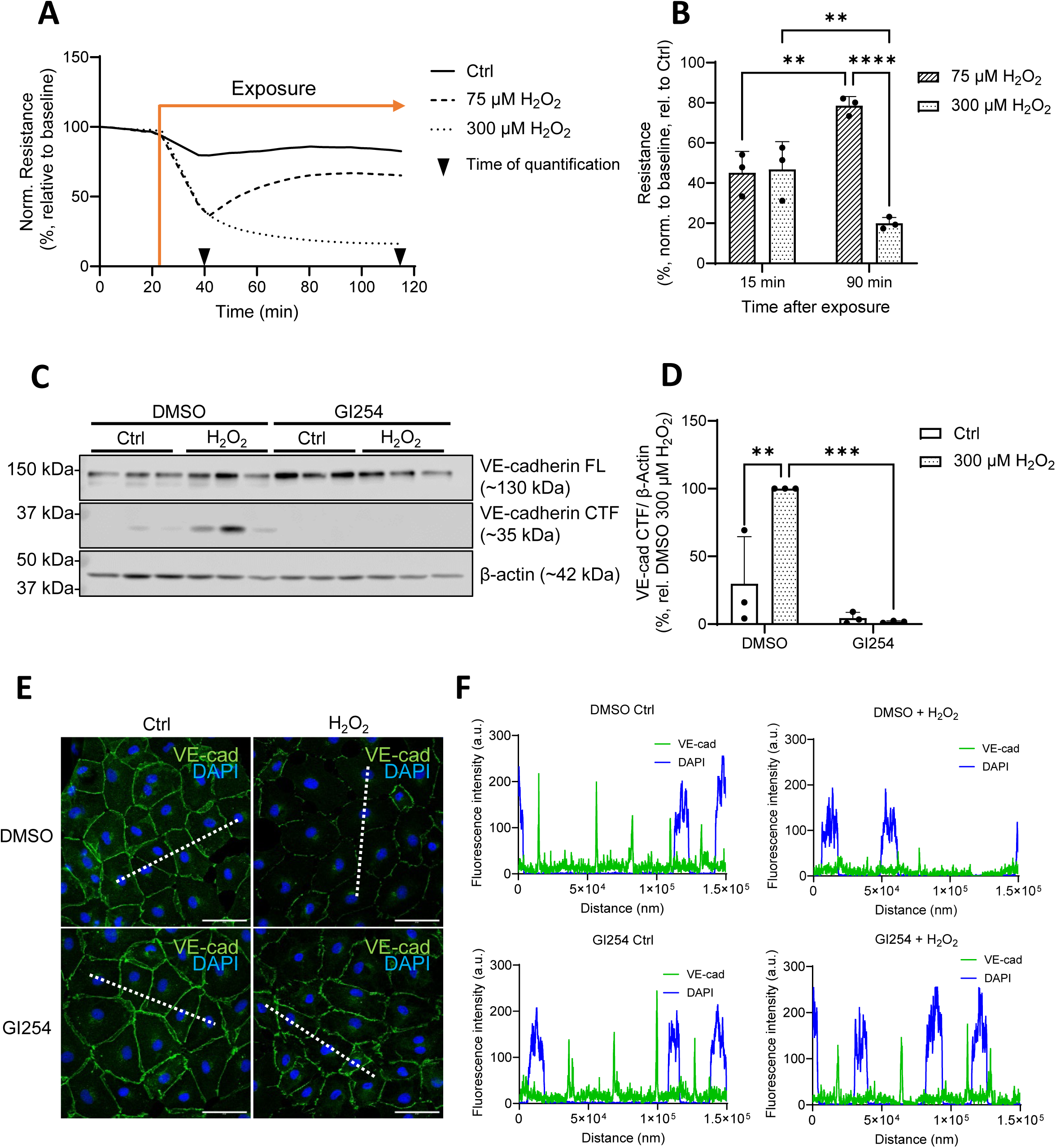
H_2_O_2_ induces HPMEC barrier instability and ADAM10-dependent VE-cadherin cleavage. Changes in HPMEC electrical resistance (normalized to baseline levels) were recorded with an ECIS device at 4000 Hz upon application of H_2_O_2_ (75 µM, 300 µM) (**A**). The normalized resistance values 15 and 90 min after exposure were quantified (**B**). Representative Western blot (from one donor, 3 technical replicates) of the full length (FL) and C-terminal fragment (CTF) levels of VE-cadherin protein in HPMECs 2 h after H_2_O_2_ exposure (300 µM) in the presence and absence of the ADAM10 inhibitor GI254023X (GI254, 3 μM) (**C**). β-actin was probed as a loading control. Normalized levels of VE-Cadherin CTF from these Western blots were quantified (**D**). (**E**) Immunofluorescence staining of HPMECs using a VE-cadherin antibody (green) and DAPI (blue), scale bar: 50 µm. Cells were fixed 2 h after H_2_O_2_ exposure in the presence and absence of GI254 (3 μM). (**F**) Fluorescence intensity profiles (measured along the dashed white lines in (**E**)) of VE-cadherin and DAPI signal. Data reflect the mean (**A**, **B**, **D**) + SD (**B**, **D**) from 3 independent donors (*n* = 3). Significance between means was analyzed using two-way ANOVA and Tukey post hoc tests (**B**, **D**); ** *p* < 0.01, *** *p* < 0.001, **** *p* < 0.0001.

### 3.2 TRPV2 mediates ADAM10-driven VE-cadherin shedding upon H_2_O_2_ exposure

We next sought to determine the involvement of the redox-sensitive TRP channels, TRPV2 and TRPM2, in H_2_O_2_-induced HPMEC barrier dysfunction. Quantitative rt-PCR confirmed that both genes were transcribed in HPMECs (Supp. Fig. S2A), and both proteins were detected in cell lysates via Western blot (Supp. Fig. S2B, C). Ca^2+^ imaging experiments revealed that the increase of intracellular Ca^2+^ ([Ca^2+^]_i_) upon H_2_O_2_ exposure was dependent on both channels (Supp. Fig S2D, E). At the protein level, pre- and co-incubation with the TRPV2 inhibitor tranilast [31] significantly reduced the H_2_O_2_-driven, ADAM10-mediated cleavage of VE-cadherin (Fig. 2A) when quantified (Fig. 2B). The TRPV2/ADAM10 VE-cadherin cleavage pathway was further confirmed with the TRPV2 activator, cannabidiol (CBD) [27] (Fig. 2C, quantified in D). Conversely, inhibition of TRPM2 with econazole [32] increased the formation of the H_2_O_2_-driven VE-cadherin cleavage product (Fig. 2E, quantified in F). These results were corroborated using alternate TRPV2 and TRPM2 inhibitors (Supp. Fig. S3A-D). Notably, while gene transcripts of the redox-sensitive TRPV4 channel were also detected in HPMECs (Supp. Fig. S2A), pretreatment with the specific TRPV4 inhibitor GSK2193874 [33] had no effect on the degree of H_2_O_2_–induced VE-cadherin CTF formation (Supp. Fig. S3E, F).

**Fig. 2:**
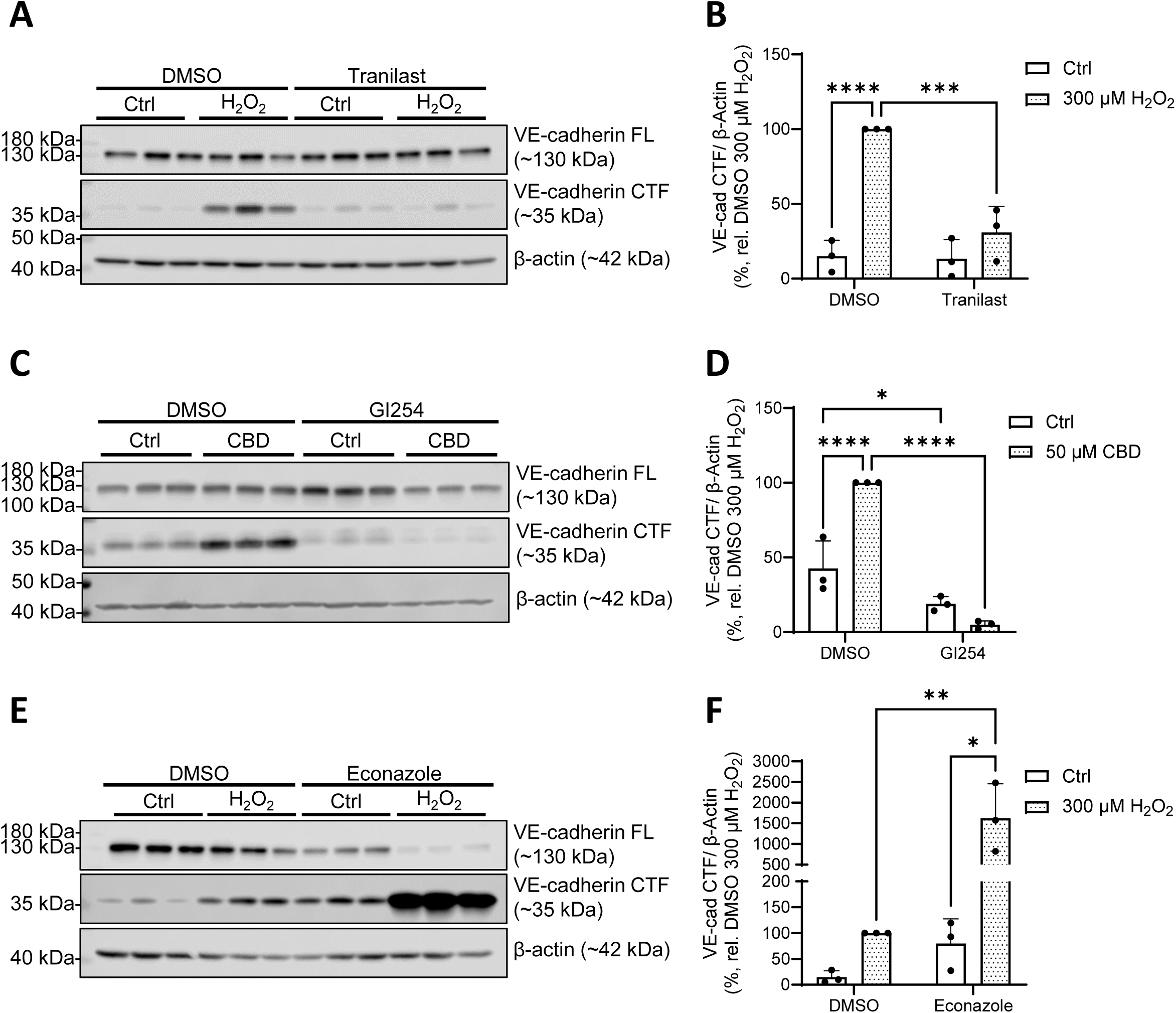
TRPV2 and TRPM2 VE-cadherin cleavage in HPMECs. Representative Western blot (from one donor, 3 technical replicates) of FL and CTF VE-cadherin protein levels in HPMECs upon TRPV2 inhibition (50 µM tranilast) and 2 h exposure to H_2_O_2_ (300 µM; **A**, quantified in **B**). (**C**) Representative Western blot (from one donor, 3 technical replicates) of FL and CTF VE-cadherin protein levels in HPMECs upon ADAM10 inhibition (3 µM GI254023X, GI254) and 2 h exposure to cannabidiol (CBD, 50 µM), quantified in (**D**). Representative Western blots (from one donor, 3 technical replicates) of FL and CTF VE-cadherin protein levels in HPMECs upon TRPM2 inhibition (10 µM econazole) and 2 h exposure to H_2_O_2_ (300 µM; **E**, quantified in **F**). β-actin was probed as a loading control. Quantified data (**B**, **D, F**) reflect the mean + SD from 3 independent donors (*n* = 3). Significance between means was analyzed using two-way ANOVA, with Tukey post hoc tests; * *p* < 0.05, ** *p* < 0.01, *** *p* < 0.001, **** *p* < 0.0001.

### 3.3. Recovery of HPMEC barrier integrity requires TRPV2 and TRPM2 functionality

We next assessed whether inhibition of TRPV2 and TRPM2 channels would influence the HPMEC barrier response to H_2_O_2_. The protective effect of TRPM2 on VE-cadherin cleavage was also evident in our measurements of barrier resistance, as application of the TRPM2 inhibitor econazole prior to the addition of 75 μM H_2_O_2_ significantly impaired HPMEC barrier recovery (Fig. 3A), with resistance values dropping to 33% ± 23% of control values after 90 minutes of treatment (Fig. 3B). In addition to TRPM2, TRPV2 activity was also necessary for HPMEC barrier recovery. HPMECs exposed to 75 μM H_2_O_2_ experienced significantly reduced recovery when pretreated with tranilast (Fig. 3C, quantified in Fig. 3D).

**Fig. 3:**
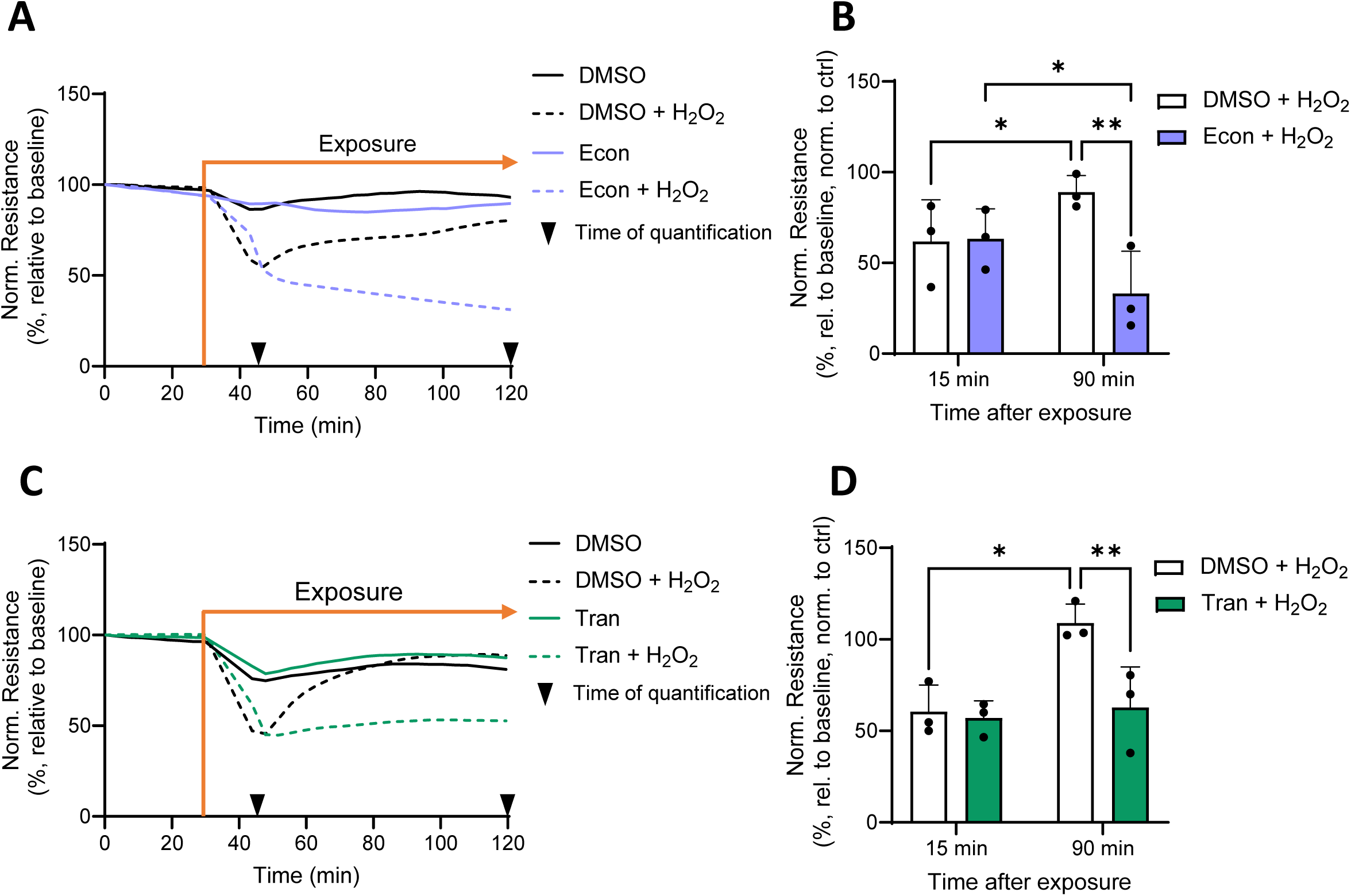
TRPM2 and TRPV2 facilitate HPMEC barrier recovery following H_2_O_2_ exposure. Changes in barrier resistance (normalized to baseline) were measured in HPMECs which were preincubated with DMSO or econazole (econ, 10 µM, 1 h) and subsequently exposed to 75 µM H_2_O_2_ (**A**). HPMEC resistance values (presented as % of control values) were quantified 15 and 90 min after H_2_O_2_ application **(B)** Similar experiments were conducted with the TRPV2 inhibitor tranilast (tran, 50 µM, 1 h preincubation, (**C, D**)). Data represent mean (**A**-**D**) + SD (**B**, **D**) of results from 3 independent donors (*n* = 3), and significance between means was analyzed with two-way ANOVA and Tukey post hoc tests; * *p* < 0.05, ** *p* < 0.01.

### 3.4. TRPV2 facilitates HPMEC barrier recovery through “cadherin switching”

TRPV2-driven VE-cadherin cleavage could facilitate HPMEC barrier recovery by enabling the translocation of neural cadherin (N-cadherin) to the plasma membrane, promoting wound healing. A time-course series of immunofluorescence stainings revealed that, upon exposure to 75 μM H_2_O_2_, VE-cadherin was rapidly removed from the cell surface after 15 minutes, but reappeared within 90 minutes (Fig. 4A). In contrast, N-cadherin, while initially dispersed in the intracellular space, organized at the plasma membrane 15 minutes after H_2_O_2_ exposure, returning to the intracellular space within 90 minutes (Fig. 4B). An analysis of fluorescence intensity profiles drawn between adjacent HPMECs (see Supp. Fig. 4A, B) confirmed that both the transient loss of VE-cadherin and the increase in N-cadherin signal at the cell borders 15 minutes after H_2_O_2_ exposure were significantly reduced upon either TRPV2 or ADAM10 inhibition (Fig. 4C, D). In HPMECs pretreated with the TRPM2 inhibitor econazole, VE-cadherin was also lost from the plasma membrane upon H_2_O_2_ application, concurrent with N-cadherin recruitment. However, VE- cadherin signal remained absent from the plasma membrane even at 90 minutes post-exposure (Supp. Fig. S4C-F).

**Fig. 4:**
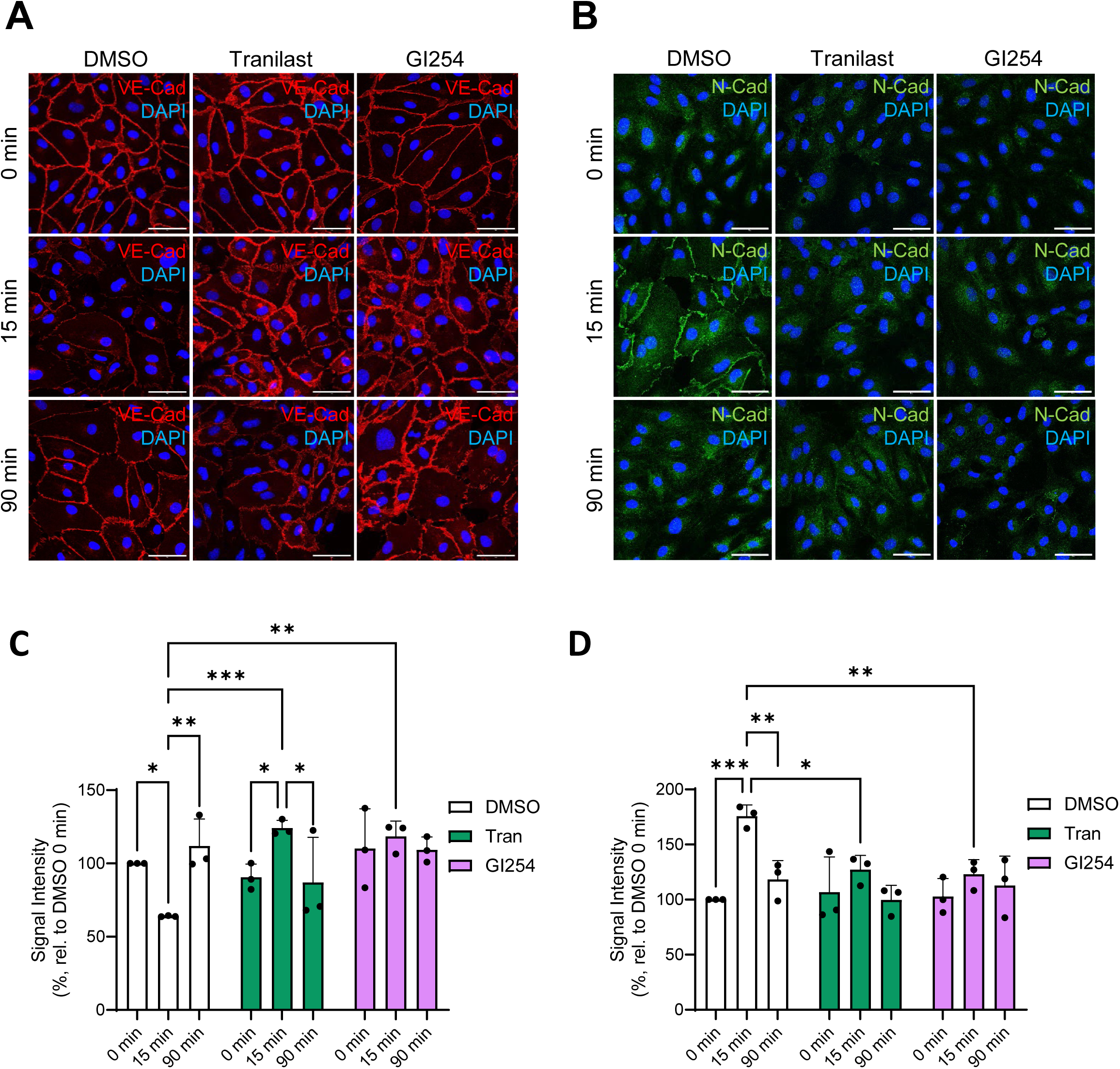
TRPV2 and ADAM10 are necessary for altered localization of N- and VE-cadherin following H_2_O_2_ exposure. (**A**) HPMEC immunofluorescence staining of VE-cadherin (red) over a timecourse of H_2_O_2_ exposure (75 µM; 0 min, 15 min, 90 min) in the presence and absence of TRPV2 and ADAM10 inhibitors (50 µM tranilast, 3 µM GI254023X, respectively). Further immunofluorescence stainings for N-cadherin (green) were conducted in the same conditions (**B**). Nuclei were stained with DAPI (blue), scale bars: 50 µm. Mean peak signal intensities of VE-cadherin (**C**) and N-cadherin (**D**) from fluorescence intensity profiles were quantified from stainings performed in HPMECs from three different donors (*n* = 3, 30 regions per *n*), and significance between means was analyzed with two-way ANOVA and Tukey post hoc tests; * *p* < 0.05, ** *p* < 0.01, *** *p* < 0.001.

## 4. Discussion

ROS are not only mediators of vascular pathology, but are also critical signaling molecules for endothelial cell proliferation, growth and motility [7,8,13,34]. In this study, we describe a pathway by which TRPV2 channels, alongside TRPM2 channels, modulate AJ protein composition and facilitate the recovery of HPMEC barrier function after H_2_O_2_ exposure.

We found that TRPV2, a redox-sensitive channel, played a significant role in mediating HPMEC response to ROS. H_2_O_2_ exposure triggered an [Ca^2+^]_i_ increase within 5 minutes, a reaction which was significantly reduced upon pharmacological inhibition of TRPV2. During oxidative signaling, TRPV2 localizes at the cell podosome, a membrane region that supports cell motility through localized proteolysis [35,36]. Along this line, we observed that TRPV2 inhibition prevented the H_2_O_2_-driven cleavage of VE-cadherin, a known substrate of the Ca^2+^-activated protease ADAM10 [17]. Therefore, we propose a TRPV2/ADAM10/VE-cadherin pathway through which HPMECs respond to ROS via ectodomain cleavage of VE-cadherin. This cleavage event may not be restricted to ADAM10 and VE-cadherin, and the involvement of other Ca^2+^-activated proteases and their cell-adhesion substrates presents a promising avenue for further study. In addition, it remains to be determined whether the extracellular fragment of VE-cadherin released during this process plays a role in downstream signaling, as is the case with its epithelial counterpart E-cadherin [37].

Our study also offers insight into the complex function of TRPM2 in mediating vascular permeability. We observed that the H_2_O_2_-driven [Ca^2+^]_i_ increase in HPMECs was significantly reduced upon pharmacological inhibition of TRPM2. These findings corroborate those of Mittal *et al.*, who also described a TRPM2-dependent increase in VE-cadherin phosphorylation at sites necessary for the protein’s internalization [22,38]. VE-cadherin internalization, which occurs upon H_2_O_2_ exposure [39], is an early step in cell migration [40] and could have significant implications for wound healing following vascular injury. We found that the return of VE-cadherin signal at the plasma membrane and recovery of monolayer integrity following H_2_O_2_ exposure were only possible with TRPM2 functionality. It is possible that, where TRPV2 activation promotes localized VE-cadherin proteolysis at the podosome, TRPM2 could facilitate temporary internalization of full-length VE-cadherin proteins until the barrier can be reestablished. This role for TRPM2 would also account for the increase in H_2_O_2_-induced VE-cadherin CTF formation upon TRPM2 inhibition.

In confluent endothelial cells, VE-cadherin localizes primarily to AJs at the plasma membrane, where it is thought to contribute to contact inhibition of cell growth and proliferation [41,42]. N-cadherin, in contrast, is associated with cell migration and wound healing and is unable to translocate to the plasma membrane in the presence of VE-cadherin at AJs [41,43]. With VE- cadherin removed from the plasma membrane, N-cadherin translocates to the cell surface, where it can form heterotypic adhesions with neighboring cells. The resulting N-cadherin adhesion complex promotes Rac1 activation, which in turn induces the recruitment of VE-cadherin back to AJs [44]. We were able to clearly observe this mechanism of cadherin-switching in our immunofluorescence stainings, which could also explain why the inhibition of the TRPV2/ADAM10/VE-cadherin cleavage pathway was detrimental to the recovery of HPMEC barrier resistance following H_2_O_2_ exposure. Our results support a pathway by which redox-sensitive TRPV2 channels facilitate the removal of VE-cadherin from HPMEC AJs through ectodomain cleavage. The resulting paracellular gaps are then rapidly repaired, possibly through N-cadherin-mediated recruitment of VE-cadherin. This pathway could be particularly relevant during leukocyte transmigration, which is characterized by H_2_O_2_ release [45], a temporary increase in paracellular permeability and a focal, transient loss of VE-cadherin complexes at AJs [4].

## CRediT authorship contributions statement

**Lena Schaller:** Conceptualization, Formal analysis, Investigation, Methodology, Visualization, Writing – original draft, Writing - review & editing. **Thomas Gudermann**: Project administration, Writing – review & editing. **Alexander Dietrich**: Conceptualization, Formal analysis, Funding acquisition, Investigation, Methodology, Project administration, Resources, Supervision, Visualization, Writing – review & editing

## Data availability

All data are available in the main text or the supplementary materials.

## Acknowledgments

The authors would like to thank Bettina Braun and Benedikt Kirmayer for their excellent technical assistance.

## Sppendix A. upplementary data

Supplementary data to this article will be found online.

## Supplementary Methods

### Reagents and Antibodies

The following reagents and antibodies were used: High glucose Dulbecco’s Modified Eagle Medium (DMEM, Thermo Fisher Scientific, Waltham, US, #41965039); hydrogen peroxide solution (Merck, Darmstadt, Germany, H1009); GI254023X (Tocris, Bristol, UK, #3995); econazole (Merck, Y0001236); JNJ-28583113 (MedChemExpress, Monmouth Junction, US, #HY-149143); tranilast (Tocris, #1098); valdecoxib (Merck, #PZ0179); cannabidiol (Cayman Chemical, #90080, Ann Arbor, MI, USA); GSK2193874 (Tocris, #5106); anti-VE-Cadherin antibody (Cell Signaling, Danvers, US, #2500); anti-N-Cadherin antibody (Cell Signaling, #13116); horseradish peroxidase (HRP)-conjugated anti-β-actin antibody (Merck, #A3854); anti-TRPM2 antibody (Bethyl, Montgomery, Texas, USA, A300-414A); anti-TRPV2 antibody (Abcam, Cambridge, UK, Ab272862); peroxidase (POX)-conjugated anti-rabbit antibody (Merck, #A1654); goat anti-rabbit IgG Alexa Fluor 488 (Thermo Fisher Scientific, #A-11008). See complete antibody information in Supp. Table S3.

### SDS-PAGE and western blot analysis

The expression of VE-cadherin protein was evaluated by Western blot analysis as previously described (*60*). Following treatment, HPMECs were lysed in 150 ul RIPA buffer (with protease and phosphatase inhibitors) for 30 min on ice. Protein concentration was quantified with the Pierce BCA Protein Assay Kit (Thermo Fisher Scientific, #23225) according to the manufacturer’s protocol. Protein samples (10 µg lysate, 1x Laemmli buffer (prepared from 5x stock: 3 ml TRIS/HCl (2.6 M), pH 6.8; 10 ml glycerin; 2 g SDS; 2 mg bromophenol blue; 5 ml β-mercaptoethanol)) were heated for 10 min at 95 °C and loaded onto an SDS-PAGE gel (4 % stacking, 10 % separating). SDS-PAGE gel electrophoresis was run for 30 min at 80 V, and then at 120 V for 90 min. Proteins were then transferred from the gel to a Roti®-PVDF membrane (Roth, Karlsruhe, Germany, #T830.1) in a wet transfer system (BioRad, Feldkirchen, Germany) at 50-60 V for 1.5 h. The membrane was then blocked with 5 % low-fat milk (Roth, #T145.2) in TBS-T (0.1 % Tween20) for 1 h at RT. All antibodies were diluted in the milk blocking solution. See Supp. Table S3 for relevant antibody information. Membranes were incubated in the primary antibody solutions overnight at 4 °C. Afterwards, membranes were washed (3 x 10 min, TBS-T) and incubated for 2 h at RT in peroxidase-conjugated secondary antibody solutions. Chemiluminescence was detected following incubation in SuperSignal West Femto or Pico maximum sensitivity substrates (Life Technologies, CA, USA, #34095 and #34580), using an Odyssey-Fc unit (Licor, Lincoln, NE, USA). Full, uncut Western blots for all experiments, donors and replicates can be accessed at https://osf.io/4uy8s/?view_only=a6c92c575798494c9c634e230bcbbb65.

### Immunocytochemistry

HPMECs were seeded on poly-L-lysine-coated 12 mm glass coverslips. After treatment, cells were washed once with cold PBS, fixed in 4 % PFA/PBS (15 min, RT), and then washed thrice with cold PBS. Cells were permeabilized for 10 min at RT in a 0.2 % Triton X-100/PBS solution, and then washed 4 x 5 min in PBS-T (0.1 % Tween20 in PBS). HPMECs were blocked for 1 h in PBS with 0.1 % Tween20 and 5 % BSA, and then incubated overnight at 4 °C in primary antibody solutions prepared in blocking buffer. See complete antibody information in Supp. Table S3. The following day, cells were washed (4 x 5 min, PBS-T), incubated for 2 h at RT in secondary antibody solutions, and washed again (4 x 5 min, PBS-T). All antibodies were diluted in blocking buffer. Nuclei were stained with DAPI (0.1 mg/l in PBS) for 3 min at RT, after which cells were washed (4 x 5 min, PBS-T). Coverslips were mounted with PermaFluor mounting medium (Epredia, Kalamazoo, MI, USA, #TA-030-FM) and kept at 4 °C. Confocal images were taken with a Zeiss LSM 880 microscope (Zeiss, Oberkochen, Germany) using the ZEN Black software (Zeiss, version 2.3). Images were processed with FIJI software (Image J v.1.53c, Wayne Rasband, NIH, USA) (*61*). Fluorescence intensity profiles were drawn in ZEN Blue software (Zeiss, version 3.4). For signal quantification, profiles were drawn between the nuclei of two adjacent HPMECs. Mean peak intensity values from ten profiles were calculated for 3 images from each condition, and these experiments were replicated thrice in different donors.

### Statistical analysis

Statistical analysis was performed with GraphPad Prism 10 software (GraphPad Software, San Diego, USA). Significant differences are indicated by asterisks, where *p* < 0.05 (*), 0.01 (**), 0.001 (***), and 0.0001 (****).

### LDH Cytotoxicity Assay

H_2_O_2_-induced cytotoxicity was assessed through an LDH assay (Merck, #11644793001), according to the manufacturer’s protocol. Briefly, cells were incubated in 300 µM H_2_O_2_ for 2h, at which point the supernatant was collected and tested for the reduction of tetrazolium salt by NADH as a measure of LDH activity. Triton X100 (2%) was applied as a positive control.

### Quantitative Reverse-Transcription (qRT)-PCR

Total RNA from HPMECs was isolated with the RNeasy Plus Mini Kit (Qiagen, Hilden, Germany, #74136). 1 µg mRNA was then transcribed to cDNA using the RevertAid H Minus First Strand cDNA Synthesis Kit (Life Technologies, Darmstadt, Germany, #K1631), with reverse transcription polymerase and random primers according to the manufacturer’s protocol. The levels of mRNA transcripts of target genes were assessed using real-time quantitative PCR, as described previously (*60*). Briefly, 3 µl of template cDNA was added to 7 µl of a master mix containing 2x Absolute QPCR SYBR Green Mix (Life Technologies, #AB1158B), 10 pmol of the respective primer pair (Metabion, Planegg, Germany, see table S4 for primer sequences) and water. For qRT-PCR, the following program was run in a light-cycler 480 device (Roche, Mannheim, Germany): activation (15 min, 94 °C); 45 cycles of denaturation (12 s, 94 °C), annealing (30 s, 50 °C) and extension (30 s, 72 °C); and melting curve analysis. The default lightcycler software (Roche, Basel, Switzerland) allowed for the calculation of crossing points (Cps), which were used to calculate gene expression values.

### Ca^2+^ imaging

HPMECs were grown on poly-L-lysine-coated 24 mm glass coverslips until 80 % confluency. On the day of measurement, HPMECs were loaded with 2 µM Fura-2-AM (Merck, #47989-1MG-F) in Ca^2+^ buffer (0.1 % BSA in HBSS (with Ca^2+^, Mg^2+^ and 0.5 M HEPES)) for 25 min at 37 °C. Coverslips were then washed with HEPES/HBSS buffer, placed into a quick-change chamber (Warner instruments, Holliston, USA, #64-0367) with 450 uL HEPES/HBSS, and positioned on the 40x oil-objective of a Leica DM98 fluorescence microscope. Changes in intracellular Ca^2+^ concentration following the application of H_2_O_2_ (300 μM, Merck, #H1009) were recorded at 340 and 380 nm wavelengths, as described (*60*). For measurements involving pharmacological inhibition, the respective inhibitors were included in both the Fura incubation solution and the treatment solutions.

### Supplementary Figure Legends

**Fig. S1.**
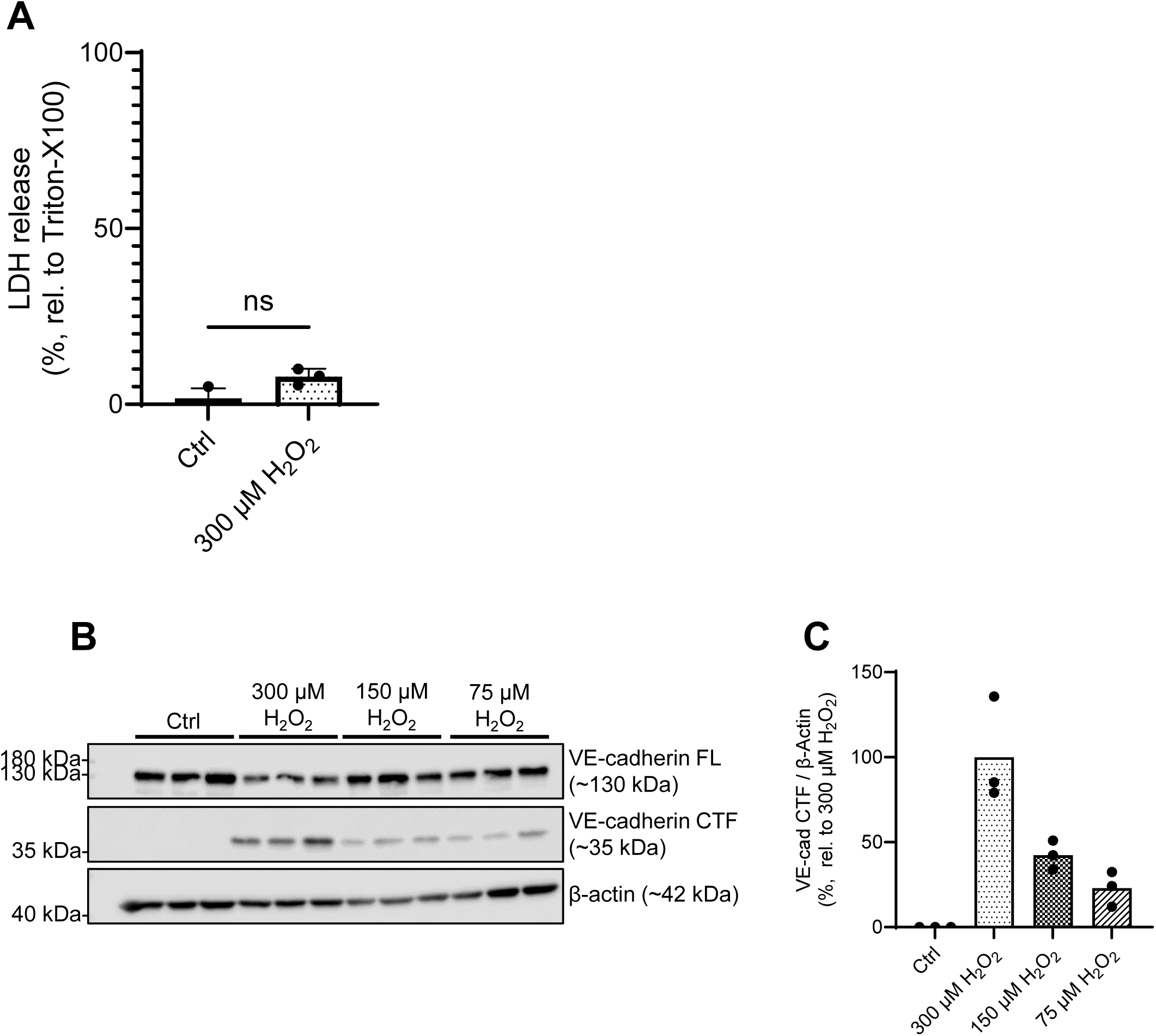
Non-cytolytic concentrations of H_2_O_2_ cause VE-cadherin degradation. (**A**) Degree of cytolysis, measured in terms of lactate dehydrogenase (LDH) activity, in HPMECs after 2 h H_2_O_2_ exposure (300 µM). Data represent the mean + SD of results from 3 independent donors (*n* = 3). Significance was assessed using a paired t test. (**B**) Western blot of the full length (FL) and C-terminal fragment (CTF) levels of VE-cadherin protein in HPMECs 2 h after H_2_O_2_ exposure at varying concentrations (75 µM, 150 µM and 300 µM). β-actin was probed as a loading control. Normalized levels of VE-Cadherin CTF from these Western blots were quantified (**C**). Data represent 3 technical replicates from one donor (*n* = 1). ns = no significance.

**Fig. S2.**
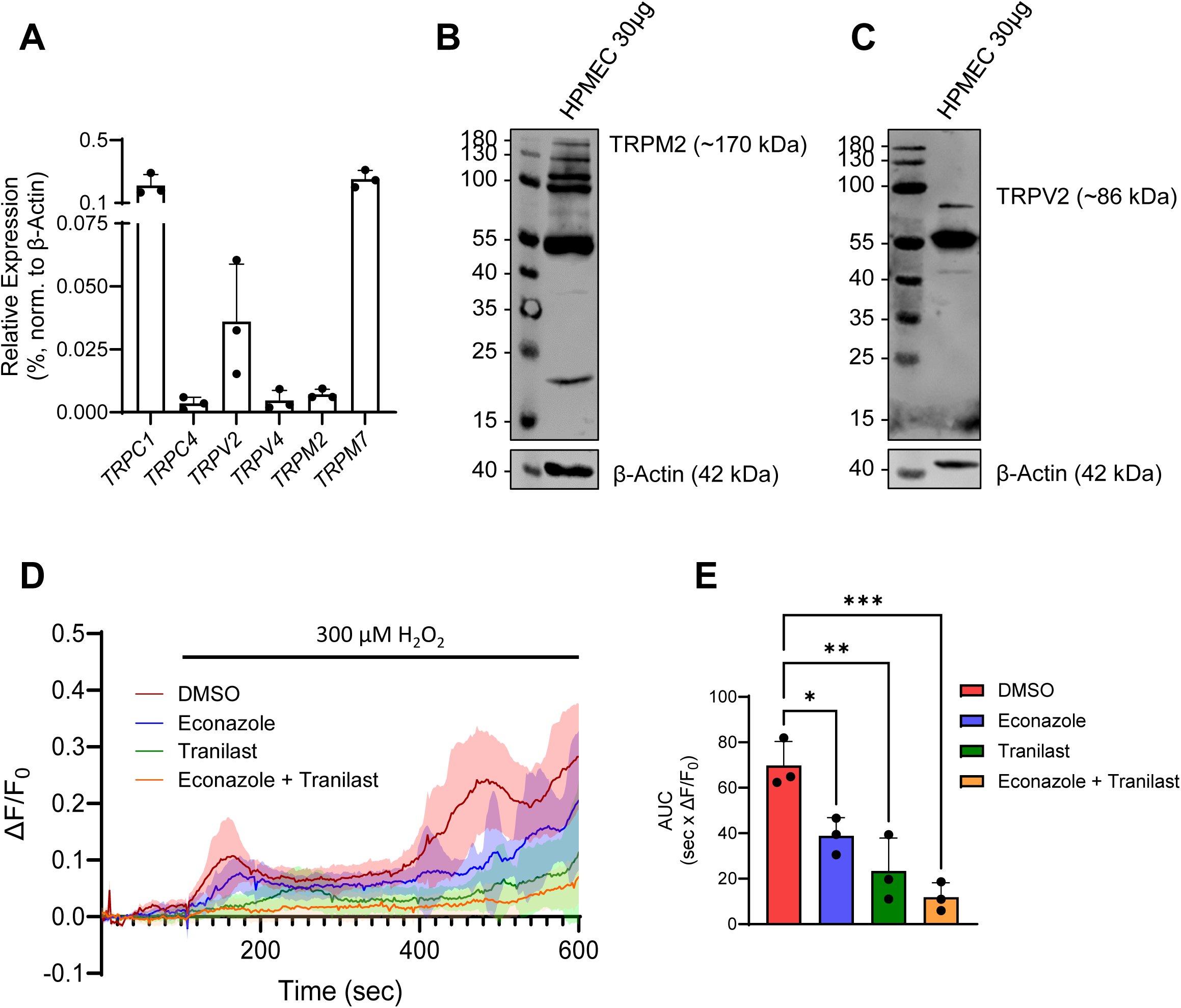
TRPM2 and TRPV2 mediate H_2_O_2_-induced Ca^2+^ influx in HPMECs. (**A**) TRP gene expression results, as detected by qRT-PCR, normalized to β-actin. Data reflect mean values + SD from 3 independent donors (*n* = 3). TRPM2 (**B**) and TRPV2 (**C**) proteins were detected in HPMEC lysates by Western blot. (**D**) Mean ΔF/F_0_ traces of HPMEC monolayer Ca^2+^ influx following H_2_O_2_ exposure (300 µM) in the presence and absence of the TRPM2 and TRPV2 inhibitors, econazole (10 µM) and tranilast (50 µM). Data represent the mean ± SD from one experiment, 35-50 cells/treatment group. This experiment was performed three times in HPMECs from a single donor at different passage numbers (*n* = 3), and the area under the curve (AUC) of each mean ΔF/F_0_ Ca^2+^ trace was quantified (**E**), with bars reflecting the mean + SEM. Significance between means was analyzed using a one-way ANOVA, with Tukey post hoc test; * *p* < 0.05, ** *p* < 0.01, *** *p* < 0.001.

**Fig. S3.**
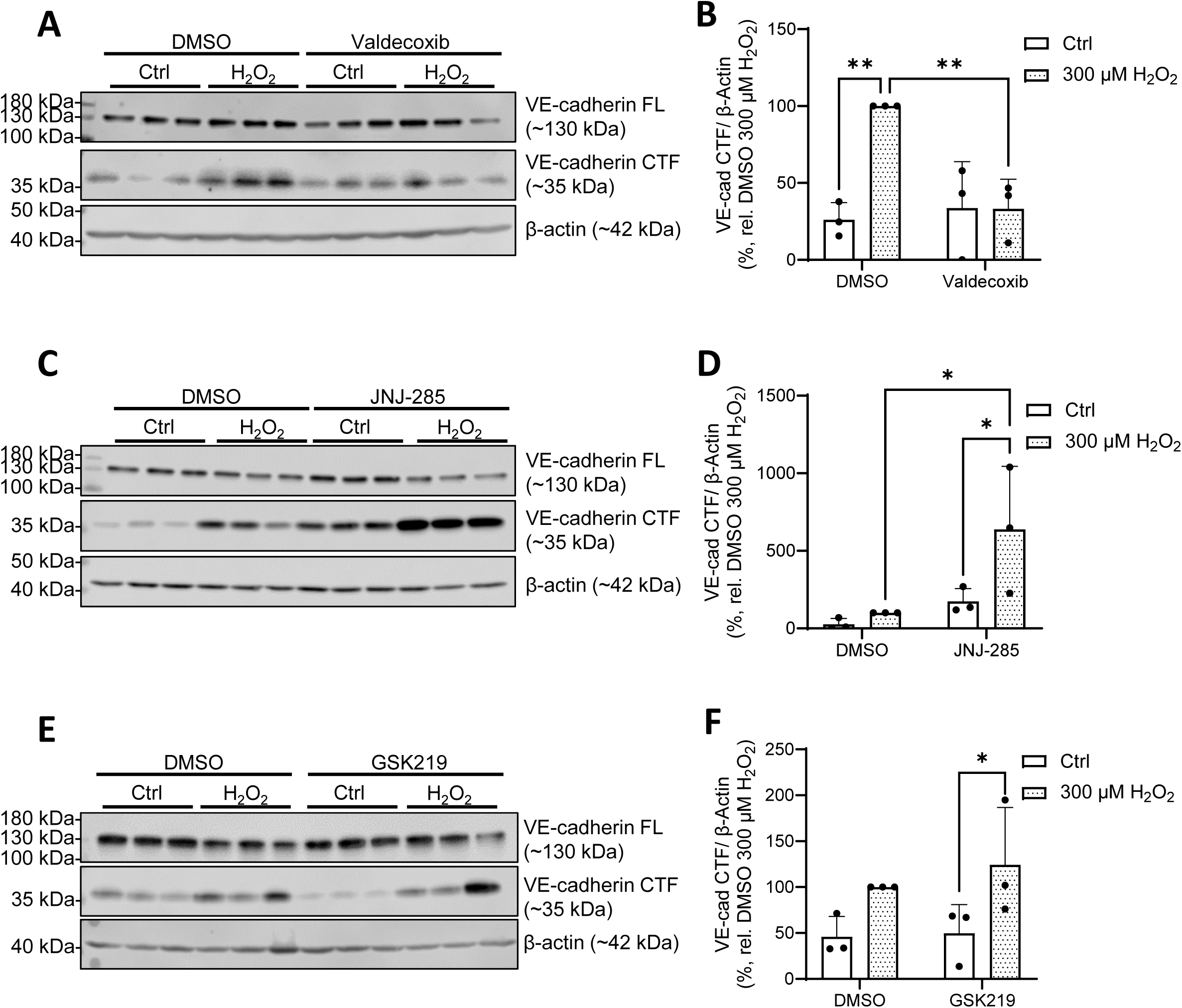
Additional controls for TRPM2 and TRPV2 modulation of HPMEC VE-cadherin upon H_2_O_2_ exposure. Representative Western blots of FL and CTF VE-cadherin protein levels after H_2_O_2_ exposure (2 h, 300 µM) upon TRPV2 (**A**, quantified in **B**), TRPM2 (**C**, quantified in **D**) and TRPV4 (**E**, quantified in **F**) inhibition (100 µM valdecoxib, 10 µM JNJ-28583113 and 300 nM GSK2193874, respectively). For all Western blots, β-actin was probed for as a loading control, samples shown are from a single donor, 3 technical replicates. Western blot quantifications represent the mean + SD of results from 3 independent donors (**B**, **D**) or 3 consecutive passages from one donor (**F**); (*n* = 3). Significance between means was analyzed using two-way ANOVA, with Tukey post hoc test; * *p* < 0.05, ** *p* < 0.01.

**Figure S4.**
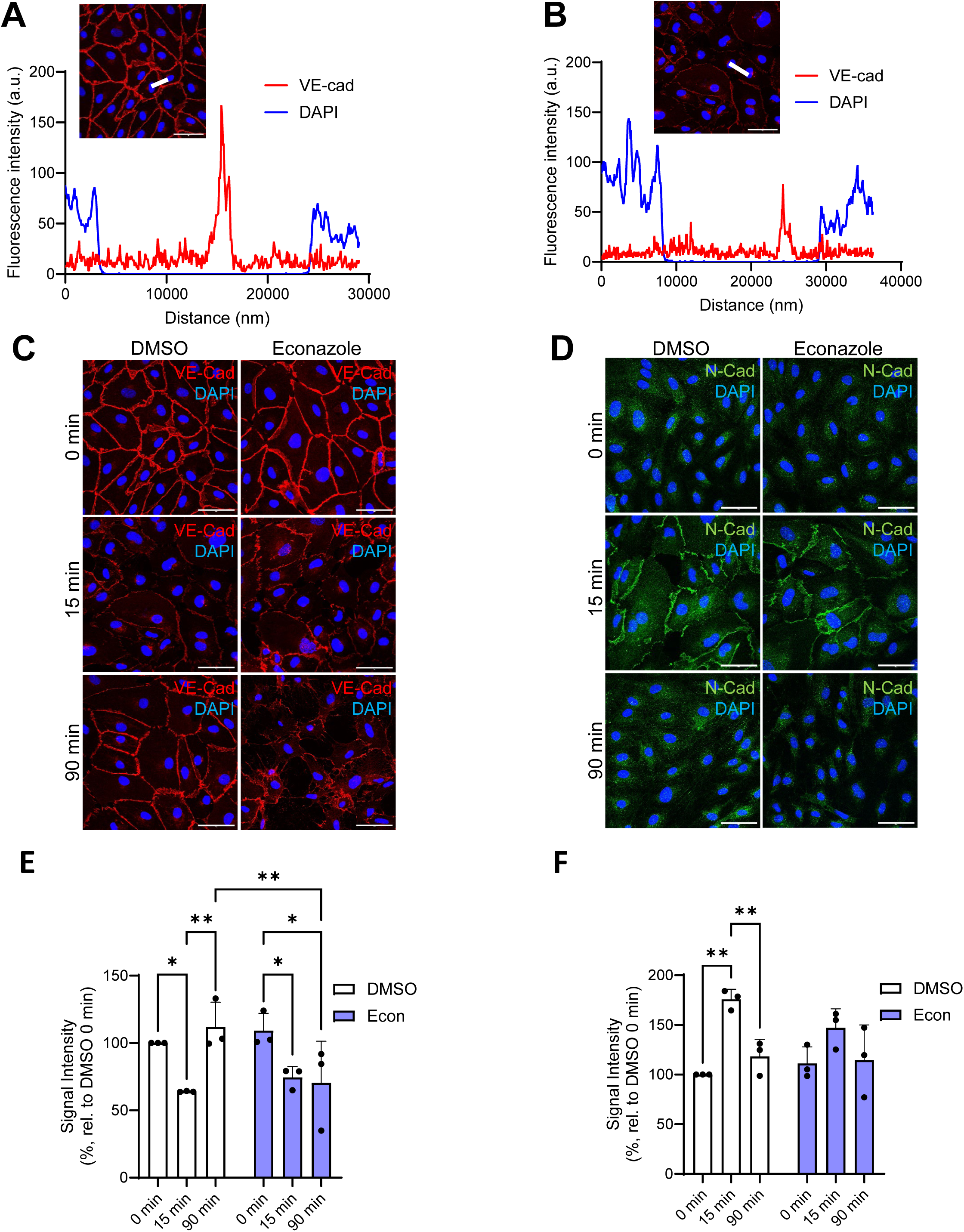
Representative fluorescence intensity profiles and immunofluorescence results for H_2_O_2_-induced changes in cadherin localization during TRP inhibition. Representative fluorescence intensity profiles (indicated by the thick white line in the image insets) for regions spanning HPMEC cell-cell junctions and adjacent nuclei in cells at 0 min (**A**) and 15 min (**B**) of H_2_O_2_ exposure (75 µM). Scale bars: 50 µm. Timecourse immunofluorescence stainings of VE-cadherin (red) (**C**) and N-cadherin (green) (**D**) over a timecourse of H_2_O_2_ exposure (75 µM; 0 min, 15 min, 90 min) in the presence and absence of the TRPM2 inhibitor econazole (10 µM). Nuclei were stained with DAPI (blue), scale bars: Mean peak signal intensities of VE-cadherin (**E**) and N-cadherin (**F**) from fluorescence intensity profiles were quantified from stainings performed in HPMECs from three different donors (*n* = 3), and significance between means was analyzed with two-way ANOVA, and Tukey post hoc tests; * *p* < 0.05, ** *p* < 0.01.

**Table S1:** HPMEC donor information (Promocell)

| Donor ID (Lot #) | Catalogue # | Age | Sex | Ethnicity | Disease Status |
| --- | --- | --- | --- | --- | --- |
| 467Z025.1 | C-12281 | 61 | Female | Caucasian | No COPD; No Asthma |
| 463Z013.1 | C-12281 | 57 | Female | Caucasian | No COPD; No Asthma |
| 489Z006.1 | C-12281 | 51 | Male | Caucasian | No COPD; No Asthma |
| 489Z005 | C-12281 | 52 | Female | Caucasian | No COPD; No Asthma |

**Table S2:**
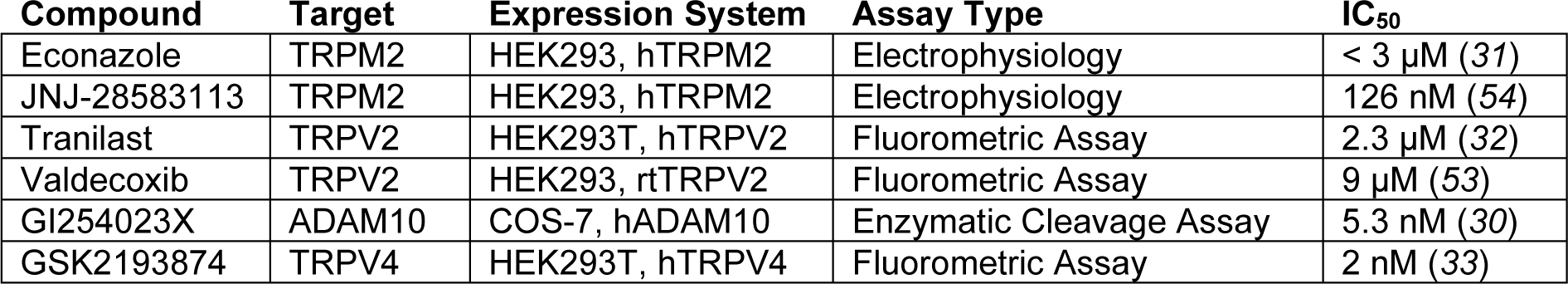
IC_50_ values for TRP and ADAM inhibitors.

**Table S3:** Antibodies used for Western blotting and immunocytochemistry.

| Primary Antibodies | Supplier | Cat. # / RRID | Dilution |
| --- | --- | --- | --- |
| VE-Cadherin (rb pAb) | Cell Signaling | 2158 / AB_2077970 | WB: 1:1000; ICC: 1:400 |
| N-Cadherin (rb pAb) | Cell Signaling | 13116 / AB_2687616 | ICC: 1:400 |
| TRPM2 (rb pAb) | Bethyl | A300-414A / AB_2208495 | WB: 1:300 |
| TRPV2 (rb pAb) | Abcam | Ab272862 / AB_2892218 | WB: 1:300 |
| B-actin-HRP (mo pAb) | Merck | A3854 / AB_262011 | WB: 1:10,000 |

| Secondary Antibodies | Supplier | Cat. # / RRID | Dilution |
| --- | --- | --- | --- |
| Mouse-HRP | Cell Signaling | 7076 / AB_330924 | WB: 1:10,000 |
| Rabbit Alexa Fluor 488 | ThermoFisher Scientific | A32731 / AB_2633280 | ICC: 1:250 |
| Mouse Alexa Fluor 594 | ThermoFisher Scientific | A-11005 / AB_2534073 | ICC: 1:250 |

**Table S4:**
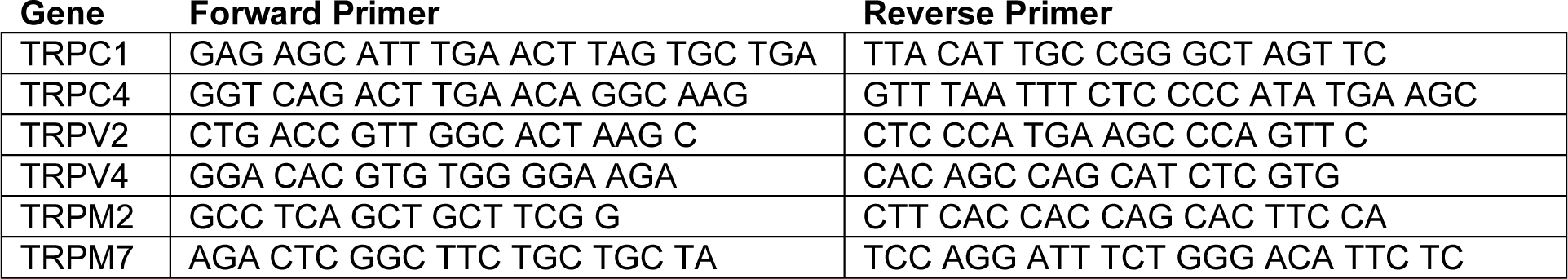
DNA-sequences of qRT-PCR primer pairs.

